# Frequent lack of repressive capacity of promoter DNA methylation identified through genome-wide epigenomic manipulation

**DOI:** 10.1101/170506

**Authors:** Ethan Ford, Matthew R. Grimmer, Sabine Stolzenburg, Ozren Bogdanovic, Alex de Mendoza, Peggy J. Farnham, Pilar Blancafort, Ryan Lister

## Abstract

It is widely assumed that the addition of DNA methylation at CpG rich gene promoters silences gene transcription. However, this conclusion is largely drawn from the observation that promoter DNA methylation inversely correlates with gene expression in natural conditions. The effect of induced DNA methylation on endogenous promoters has yet to be comprehensively assessed. Here, we induced the simultaneous methylation of thousands of promoters in the genome of human cells using an engineered zinc finger-DNMT3A fusion protein, enabling assessment of the effect of forced DNA methylation upon transcription, histone modifications, and DNA methylation persistence after the removal of the fusion protein. We find that DNA methylation is frequently insufficient to transcriptionally repress promoters. Furthermore, DNA methylation deposited at promoter regions associated with H3K4me3 is rapidly erased after removal of the zinc finger-DNMT3A fusion protein. Finally, we demonstrate that induced DNA methylation can exist simultaneously on promoter nucleosomes that possess the active histone modification H3K4me3, or DNA bound by the initiated form of RNA polymerase II. These findings suggest that promoter DNA methylation is not generally sufficient for transcriptional inactivation, with implications for the emerging field of epigenome engineering.

**One Sentence Summary:** Genome-wide epigenomic manipulation of thousands of human promoters reveals that induced promoter DNA methylation is unstable and frequently does not function as a primary instructive biochemical signal for gene silencing and chromatin reconfiguration.

## Main Text

While the mammalian genome is almost entirely methylated at cytosines in the CpG dinucleotide context (mCG) (*1*), regulatory regions such as CpG rich promoters and active enhancers generally exhibit a low- or un-methylated state (*2*, *3*). However, a broader understanding of whether DNA methylation at such regions is generally causative for transcriptional silencing has remained elusive. DNA methylation of gene promoters is frequently inversely correlated with transcriptional activity (*2*, *4*, *5*), and abolition of DNA methyltransferase activity through chemical inhibition or genetic disruption causes global demethylation and activates numerous genes (*6*, *7*). Such observations have led to the common conclusion that DNA methylation of CpG island (CGI) promoters and other regulatory sequences causes transcriptional silencing. However, these observations are correlative and challenging to interpret because genome-wide demethylation could have widespread and complex downstream effects upon chromatin structure and gene expression. Transfection and microinjection of differentially methylated sequences have demonstrated a repressive role of DNA methylation, but may not accurately reproduce the chromatin state and other regulatory processes of endogenous promoters (*8*–*10*). Moreover, DNA methylation has been reported to occur downstream of transcriptional regulation in some contexts. In early development and germline cells, transcription can occur from genes with methylated promoters (*11*–*13*). Furthermore, transcriptional silencing can precede the acquisition of promoter DNA methylation (*14*), including genes on the female inactive X chromosome (*15*, *16*) and imprinted genes during development (*17*). Finally, while aberrant promoter CGI hypermethylation is observed at many silenced genes in cancers, these genes are frequently already repressed in normal preneoplastic cells (*18*, *19*). Thus, whether DNA methylation at promoters generally functions as a primary instructive biochemical signal for gene silencing remains unresolved.

Targeted methods to alter methylation state at endogenous loci have been developed in recent years, including customized zinc finger (ZF) domains fused to the catalytic domain of human DNA methyltransferase 3A (ZF-DNMT3A) (*20*–*25*) or the bacterial methyltransferase M.SssI and more recently the fusion of DNMT3A to transcription activator-like effector (TALE) proteins or nuclease inactive Cas9 (*27*–*32*). These artificial epigenome modifiers allow targeted interrogation of whether DNA methylation correlates with, or is causative for, transcriptional repression. Their application at a few tested loci has indicated that induction of methylation on promoters is sometimes sufficient to repress transcription. Indeed, DNMT3B-mediated targeted induction of DNA methylation at an artificial promoter and reporter locus in mammalian (CHO)-K1 cells was sufficient to cause stable transgene silencing (*33*). However, a broad examination of the capacity of induced DNA methylation of endogenous promoters in the mammalian genome to cause changes in gene expression and chromatin has not previously been undertaken.

Given the efficient post-replicative maintenance of DNA methylation patterns by DNMT1 (*34*), DNA methylation has been implicated in long-term gene silencing (*35*). However, the stability of targeted induced methylation appears to be variable, with examples of both stable (*36*) and transient (*24*, *30*) methylation and silencing of different genes. Thus, the context in which DNA methylation is sufficient to establish transcriptional repression and the stability of induced DNA methylation has not been broadly characterized. The function of changes in DNA methylation observed in differentiation, development and disease are very challenging to interpret without a clear determination of whether the addition of DNA methylation to diverse regulatory regions is sufficient to induce transcriptional silencing. To address this, we undertook a broad assessment of the consequences upon gene expression and chromatin state of inducing DNA methylation at promoter regions throughout the human genome.

## Genome-wide epigenomic manipulation

We took advantage of a cell line derived from human breast adenocarcinoma MCF-7 cells that, upon doxycycline treatment, expresses an artificial epigenome modifier composed of 6 zinc finger (ZF) domains linked to the catalytic domain of human DNMT3A (*25*) (ZF-D3A, Fig. 1A). The ZF array was designed to bind to a GC-rich 18 bp sequence in a CGI in the *SOX2* promoter, as confirmed by ChIP-seq of the HA-epitope-tagged ZF-D3A protein (Fig. 1B). To assess the effect of ZF-D3A expression upon DNA methylation states throughout the genome, we performed whole genome bisulfite sequencing (WGBS) of 1) control MCF-7 cells harboring an integrated empty vector construct and grown in the presence of doxycycline for 3 days (MCF7-control) to establish baseline methylation levels, 2) MCF7-ZF-D3A cells induced with doxycycline for 3 days (ZF-D3A +dox), and 3) MCF7-ZF-D3A cells induced with doxycycline for 3 days then subsequently grown in doxycycline-free media for 9 days after doxycycline withdrawal (ZF-D3A dox-wd). The weighted CG methylation level (see methods) of the *SOX2* promoter was 34% higher in ZF-D3A +dox compared to MCF7-control, and approximately half of the induced methylation remained in ZF-D3A dox-wd cells (Fig. 1B). RNA-seq analysis revealed that the induction of methylation was associated with a 2.5 fold decrease in *SOX2* mRNA relative to MCF7-control cells, which was sustained after doxycycline withdrawal (Fig. 1B). Thus, ZF-D3A methylates its target region and results in repression of the *SOX2* gene.

**Fig. 1.**
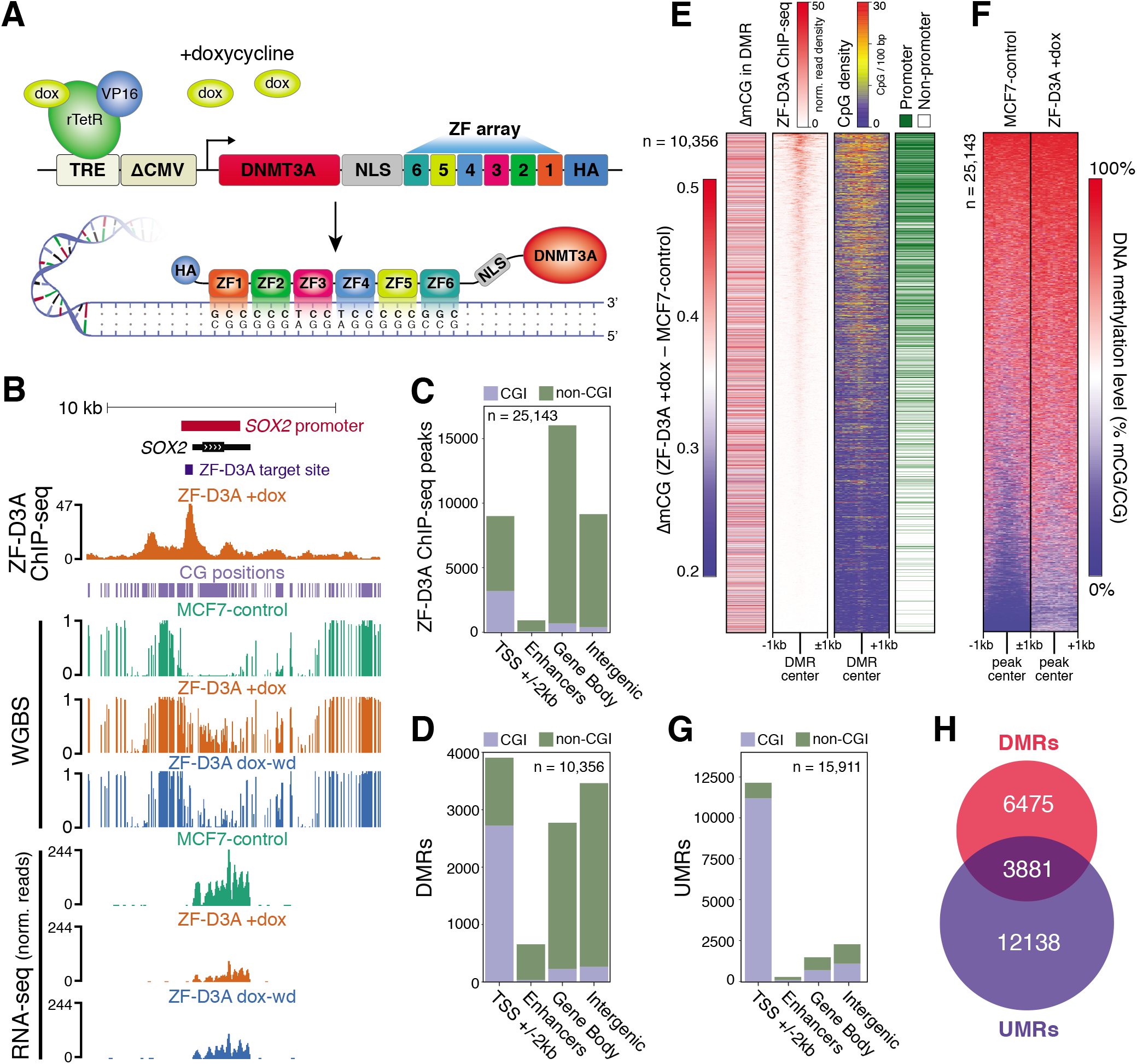
Genome-wide manipulation DNA methylation with ZF-D3A. **(A)** Schematic representation of the ZF-D3A doxycycline inducible system. **(B)** Genome browser representation of the *SOX2* locus. The promoter is defined as the unmethylated region spanning the *SOX2* transcriptional start site in MCF7-control. Genomic distribution of **(C)** identified ZF-D3A binding sites, and **(D)** differentially methylated regions (DMRs) between MCF7-control and ZF-D3A +dox. **(E)** Shown for all DMRs identified between ZF-D3A +dox and MCF7-control is: ∆mCG (difference in the methylation level [ratio of C base calls to total base calls for all CG dinucleotides in region, mCG/CG] between ZF-D3A +dox and MCF7-control), ZF-D3A ChIP-seq signal (normalized ChIP-seq read density, ±1kb flanking DMR center), CpG dinucleotide density (CpG/100bp, ±1kb flanking DMR center), and classification of each DMR as promoter or non-promoter located. **(F)** DNA methylation level in MCF7-control and ZF-D3A +dox cells flanking (±1 kb, 100 bp bins) the center of ZF-D3A ChIP-seq peaks identified in ZF-D3A +dox. **(G)** Genomic distribution of unmethylated regions (UMRs) in MCF7-control. **(H)** DMRs and UMRs with an intersection of ≥1 bp.

However, ZF domains have a propensity to bind to sequences similar to their intended target (*37*), and to GC-rich sequences in general (*38*). Given the GC-rich characteristics of the intended target site within a CGI in the *SOX2* promoter, and the prevalence of CpG-rich regulatory regions (e.g. CGIs, gene promoters), the ZF-D3A protein could potentially bind to and deposit DNA methylation broadly at CpG-rich regulatory regions throughout the genome. Indeed, analysis of the localization of HA-tagged ZF-D3A protein by ChIP-seq in ZF-D3A +dox cells identified 25,142 off-target binding sites throughout the genome (FDR <0.05), primarily in promoters, gene bodies, and intergenic regions (Fig. 1C, table S1). Therefore, this system enables the effect of induced DNA methylation to be assessed at many thousands of regulatory regions simultaneously, rather than laboriously testing one at a time.

Global methylation levels in the CH (H = A, C or T) sequence context in both MCF7-control and ZF-D3A +dox cells was equal to the bisulfite non-conversion rate (0.5%), indicating that the ZF-D3A fusion protein did not significantly alter CH methylation in the genome. In contrast, global CG methylation levels were 5.2% higher in ZF-D3A +dox cells (68.2%) compared to MCF7-control (63%). Consequently, further analyses focused on CG methylation. We identified 10,356 differentially methylated regions (DMRs) in the CG dinucleotide context between MCF7-control and ZF-D3A +dox, located in gene promoters (38%; ±2 kb of a transcriptional start site), gene bodies (27%), putative enhancers (6%; H3K4me1/H3K27ac modified chromatin), and intergenic regions not corresponding to enhancers (30%; Fig. 1D, table S2). Notably, only 35% of DMRs intersected with ZF-D3A ChIP-seq peaks. While there was an enrichment of ZF-D3A binding in the majority of DMRs, particularly those in CpG-rich promoters, many DMRs showed no detectable interaction with ZF-D3A (Fig. 1E), suggesting that ZF-D3A may induce methylation in the absence of apparent strong and persistent protein binding. The binding sites of the ZF-D3A fusion protein in ZF-D3A +dox cells exhibited a wide range of initial methylation states in MCF7-control cells, with unmethylated and partially methylated regions frequently gaining methylation when the fusion protein was expressed in ZF-D3A +dox cells (Fig. 1F).

In addition to DMR detection, we defined the boundaries of unmethylated regions (UMRs, (*39*)) of the genome flanked by methylated DNA in MCF7-control cells, which frequently encompass gene promoters. This enables measurement of the DNA methylation induced across an entire gene promoter, as opposed to potentially only a subset of it, as is possible with DMRs. In MCF7-control cells 15,911 UMRs were identified (see methods), with most (77%) within 2 kb of a transcriptional start site and 82% overlapping with CGIs (Fig. 1G, table S3). UMRs and DMRs frequently overlapped, with approximately one third of each type of region intersecting with the other (Fig. 1H). Both UMRs and DMRs were used to assess the effects of the widespread binding and induction of DNA methylation by ZF-D3A.

The level and stability of DNA methylation induced by ZF-D3A was assessed by comparing the methylation level in each DMR (mCG/CG: the ratio of C base calls to total base calls for all CG dinucleotides in the region) between MCF7-control, ZF-D3A +dox, and ZF-D3A dox-wd cells (Fig. 2, A to C). The DNA methylation level was higher in ZF-D3A +dox cells compared to MCF7-control in 99.9% of DMRs, while widespread loss of the induced methylation in the DMRs was observed in ZF-D3A dox-wd (fig. S1A). To assess whether retention of induced DNA methylation in DMRs correlated with the baseline level of methylation in the absence of ZF-D3A, we focussed on DMRs with either low (<0.1) or high (>0.3) initial methylation levels in MCF7-control cells. After doxycycline induction and subsequent withdrawal in ZF-D3A cells, 66% of DMRs that had low initial methylation levels returned to baseline levels after doxycycline withdrawal (∆mCG <0.1, where ∆mCG = difference in methylation level between ZF-D3A dox-wd and MCF7-control; Fig. 2, A and C, fig. S1A). In contrast, only 8% of DMRs with high initial DNA methylation levels returned to their initial levels (∆mCG <0.1) after doxycycline withdrawal. DMRs that lost induced methylation tended to be CpG-rich and located at promoters, whereas induced DNA methylation was generally more stable at CpG-poor non-promoter genomic regions that were initially partially methylated (fig. S1A). Indeed, assessment of UMRs showed that upon expression of ZF-D3A, 68% of UMRs gained methylation (∆mCG >0.1), however nearly all UMRs lost the induced DNA methylation in the ZF-D3A dox-wd cells (Fig. 2, D to F). Therefore, UMRs are inherently resistant to the stable maintenance of artificially induced DNA methylation.

**Fig. 2.**
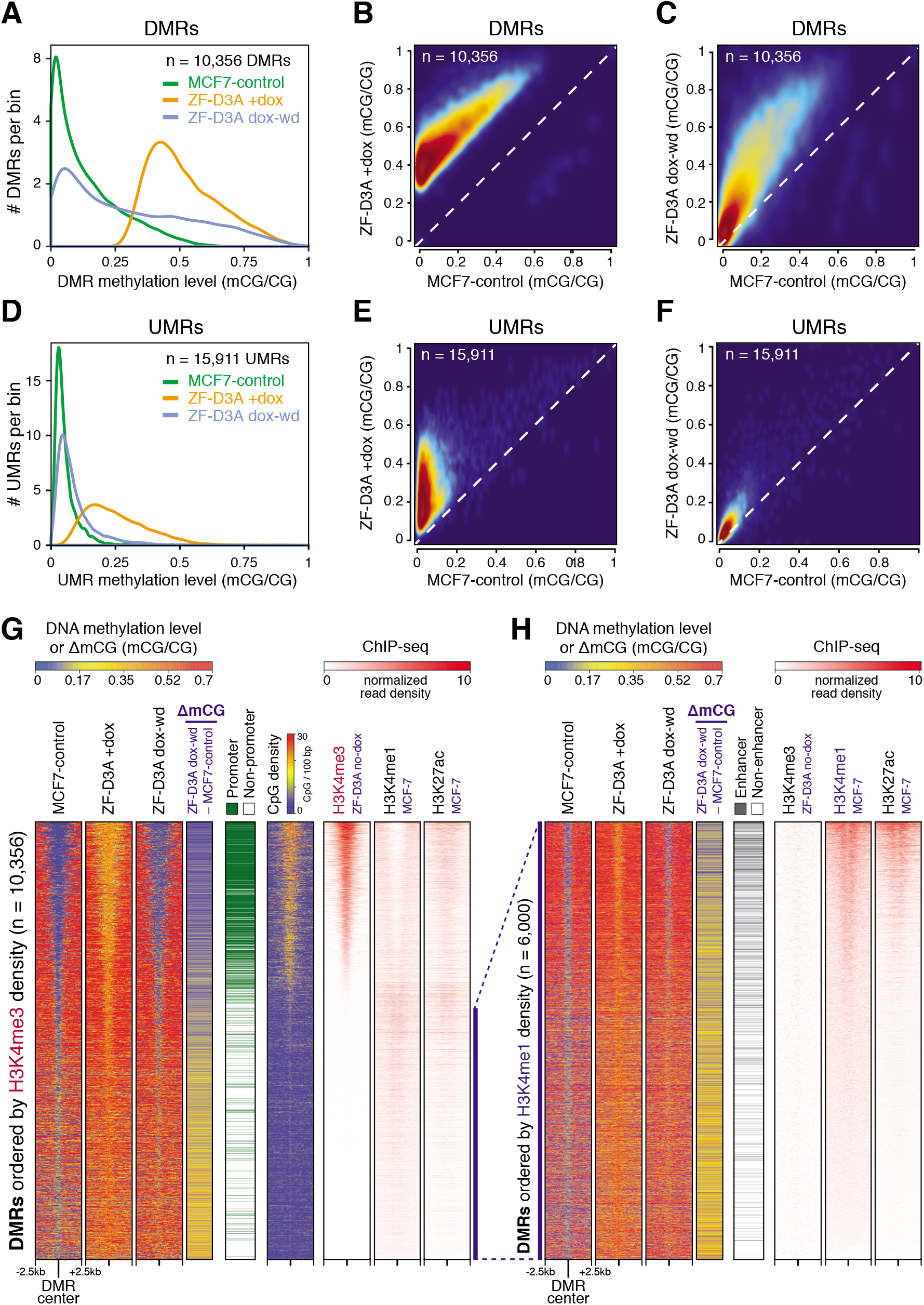
Induced methylation is frequently transient and preferentially lost from H3K4me3 marked regions. **(A)** Line histogram of DMR methylation levels in each sample. Kernel density estimate smoothed scatterplot comparison of DMR methylation levels in MCF7-control and **(B)** ZF-D3A +dox, or **(C)** ZF-D3A dox-wd. **(D)** Line histogram of UMR methylation levels in each sample. Scatterplot comparison of UMR methylation levels in MCF7-control and **(E)** ZF-D3A +dox, or **(F)** ZF-D3A dox-wd. **(G)** Heat-maps of DMRs (ordered by decreasing H3K4me3 read density) showing: DNA methylation levels (±2.5 kb), differences (ΔmCG), classification as promoter or non-promoter located, CpG dinucleotide density (±2.5 kb), and histone modification ChIP-seq read density (±2.5 kb); **(H)** As in **(G)** but for the 6,000 DMRs with the lowest H3K4me3 levels, ordered by decreasing H3K4me1 read density, and DMR classification as enhancer or non-enhancer in MCF7 (ENCODE).

As most UMRs are located in promoter regions and at CGIs (Fig. 1G), we assessed the association of all DMRs with different active chromatin states to determine whether particular histone modifications and functional genomic regions were associated with the stable or transitory nature of induced DNA methylation. ChIP-seq of H3K4me3 in ZF-D3A no-dox cells revealed that DMRs enriched for H3K4me3, which are also CpG-rich and located in promoters, are largely demethylated to baseline levels upon doxycycline withdrawal, whereas DMRs not enriched for H3K4me3 exhibit higher retention of DNA methylation (Fig. 2G). Specifically, 78% of the top quartile of DMRs, sorted by the highest H3K4me3 signal in ZF-D3A no-dox cells, returned to similar methylation levels (∆mCG <0.1) in ZF-D3A dox-wd. In contrast, only 14% of the bottom quartile of H3K4me3 sorted DMRs returned to initial methylation levels (∆mCG <0.1) after doxycycline withdrawal.

To determine whether induced DNA methylation was also transitory at regions marked with other active histone modifications, such as H3K4me1 and H3K27ac that are enriched at enhancers, we ranked the DMRs not associated with H3K4me3 by their level of H3K4me1 enrichment (Fig. 2H). The association between H3K4me1 enrichment and loss of induced DNA methylation was far less pronounced than for H3K4me3. Specifically, 9.6% of DMRs in the top quartile of H3K4me1 enrichment returned to near-baseline levels of DNA methylation (∆mCG <0.1) in ZF-D3A dox-wd, compared to 3.2% in the bottom quartile. Thus, artificially induced methylation exhibits higher stability at H3K4me1 modified loci.

## Forcibly methylated DNA associates with active chromatin marks and transcription

DNA methylation and H3K4me3 are, for the most part, mutually exclusive marks in CpG-rich regions (*40*), and H3K4me3 may protect against cytosine methylation (*41*). However, it is not known whether forced mC deposition is sufficient to induce the loss of H3K4me3. Analysis of H3K4me3 ChIP-seq read density between ZF-D3A cells not treated with doxycycline (ZF-D3A no-dox) and ZF-D3A +dox cells, at genomic regions flanking all 10,356 DMRs with a ∆mCG >0.2, revealed a small decrease (median 0.7-fold) in H3K4me3 signal after induction of ZF-D3A (Fig. 3, A and B, fig. S1B). To account for variations in ChIP efficiency, the two ChIP-seq data sets were normalized by the number of reads mapped to H3K4me3 peaks that had a ∆mCG <0.2 between ZF-D3A no-dox and ZF-D3A +dox cells (Fig. 3A, bottom). While DMRs only exhibit a ∆mCG of >0.2 upon doxycycline induction, UMRs display a broad range of induced methylation, with many that are not methylated by ZF-D3A, therefore we assessed the correlation between H3K4me3 level and DNA methylation in UMRs. Notably, there is no correlation between the change in DNA methylation and the level of H3K4me3 in UMRs (Fig. 3C). This suggests that the small decrease of H3K4me3 occupancy observed in ZF-D3A +dox cells at DMRs is not a result of induced DNA methylation, indicating that DNA methylation has minimal, if any, effect on H3K4me3 modification.

**Fig. 3.**
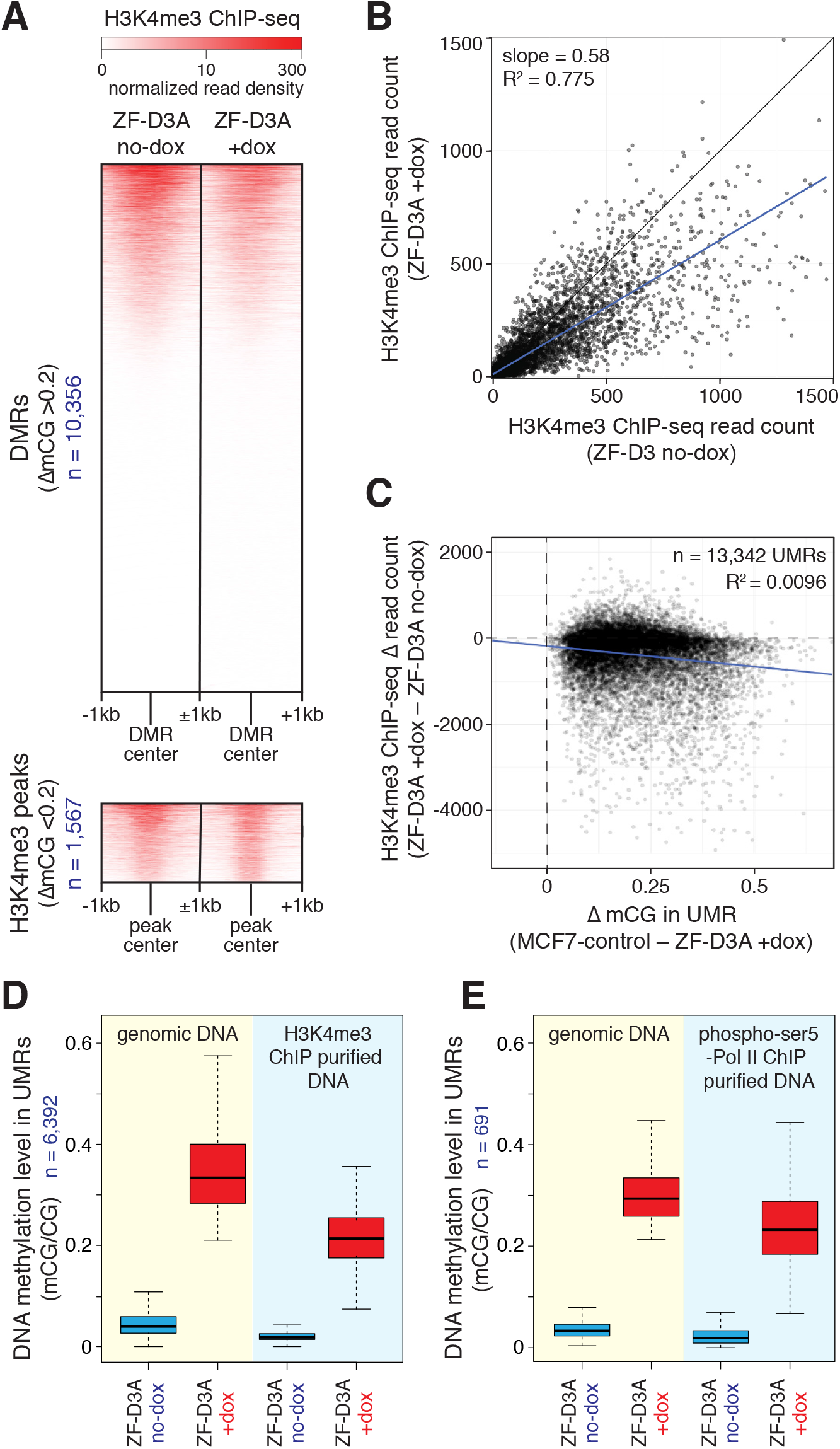
DNA methylated by ZF-D3A is bound by H3K4me3 and initiated RNA polymerase II. **(A)** H3K4me3 ChIP-seq read density flanking (±1 kb, 50 bp bins) the centre of DMRs with ∆mCG >0.2 (top), and H3K4me3 ChIP-seq peaks with ∆mCG <0.2 (bottom). **(B)** H3K4me3 ChIP-seq normalized read counts in DMRs with ∆mCG >0.2, from H3K4me3 ChIP-seq in ZF-D3A no-dox and ZF-D3A +dox cells. Black line represents equal normalized read counts in ZF-D3A no-dox and ZF-D3A +dox cells; blue line represents a linear regression of the data. **(C)** Comparison of UMR ∆mCG and difference in H3K4me3 ChIP-seq normalized read counts between ZF-D3A no-dox and ZF-D3A +dox. Blue line represents a linear regression of the data. **(D)** DNA methylation levels of UMRs that intersect with H3K4me3 ChIP-seq peaks in ZF-D3A no-dox and ZF-D3A +dox cells, for genomic DNA or DNA immunoprecipitated with an anti-H3K4me3 antibody. **(E)** DNA methylation levels of UMRs that intersect with phos-pho-ser5 RNA polymerase II ChIP-seq peaks in ZF-D3A no-dox and ZF-D3A +dox cells, for genomic DNA or DNA immunoprecipitated with an anti-phospho-ser5 RNA polymerase II antibody.

Because of the limited sensitivity of ChIP-seq and the possibility that there could be a heterogeneous population of loci with methylated and unmethylated DNA molecules, we directly measured the DNA methylation level in H3K4me3-modified chromatin using ChIP-bisulfite-sequencing (*42*–*44*). For this analysis UMRs were favored over DMRs, since UMRs span the entire unmethylated promoter region. We identified 6,392 H3K4me3 ChIP-seq peaks from both ZF-D3A +dox and ZF-D3A no-dox cells that overlapped with UMRs exhibiting a ∆mCG >0.2 across the entire UMR upon induction of the ZF-D3A fusion protein. In DNA purified by H3K4me3 ChIP in ZF-D3A +dox, 21% of CpGs within the 6,392 H3K4me3 peaks were methylated, demonstrating that H3K4me3 and DNA methylation can exist simultaneously at the same site upon forced induction of DNA methylation (Fig. 3D).

Promoter DNA methylation has been proposed to inhibit transcription, thus we investigated whether the initiated form of RNA polymerase II (phosphorylated at Serine 5 of the CTD; phospho-Ser5) was able to bind DNA methylated by ZF-D3A, by ChIP-bisulfite-sequencing (*12*) of genomic DNA isolated with an anti-phospho-Ser5 RNA polymerase II antibody. We analyzed UMRs that exhibited a ∆mCG >0.2 between ZF-D3A no-dox and ZF-D3A +dox cells and that intersected with phospho-Ser5 RNA pol II ChIP-seq peaks. Similar to H3K4me3, there was only a small decrease in the median DNA methylation level between DNA bound by phospho-Ser5 RNA pol II (29%) and non-immunoprecipitated bulk genomic DNA (23%). Importantly, phospho-Ser5 RNA pol II was clearly able to directly interact with ZF-D3A methylated DNA (Fig. 3E). Together, these data demonstrate that forced DNA methylation is not sufficient to disrupt H3K4me3 occupancy or the interaction of initiated RNA polymerase II with genomic DNA.

## Induced DNA methylation is is actively demethylated

To gain insights into the durability of induced DNA methylation after removal of ZF-D3A, DNA methylation and HA-tagged ZF-D3A protein levels were quantitated throughout doxycycline induction and subsequent withdrawal. ZF-D3A protein was strongly expressed 1 day after doxycycline addition and returned to undetectable/baseline levels 1 day after doxycycline withdrawal (Fig. 4A). The band visible at the same molecular weight as ZF-D3A prior to induction (Fig. 4A) is a non-specific protein, as indicated by the extremely similar *DNMT3A* mRNA levels in MCF7-control and ZF-D3A cells in the absence of doxycycline, and negligible *ZF* mRNA in ZF-D3A cells without doxycycline and following its withdrawal (fig. S2A). Moreover, any potential basal expression of ZF-D3A had no discernible effect on DNA methylation levels, as revealed by the very similar DMR methylation levels between MCF7-control and ZF-D3A cells without doxycycline (fig. S2B). Furthermore, PCA of RNA-seq data from MCF7-control, ZF-D3A no-dox and ZF-D3A +dox cells showed that most variance is explained by PC1 (83%), and this component clearly separates ZF-D3A +dox cells from MCF7-control and ZF-D3A no-dox, while PC2 only explains 10% of the variance and differentiates between MCF-control and ZF-D3A no-dox. Thus, MCF7-control and ZF-D3A no-dox cells are much more similar to one another than to ZF-D3A +dox cells (fig. S2C).

**Fig. 4.**
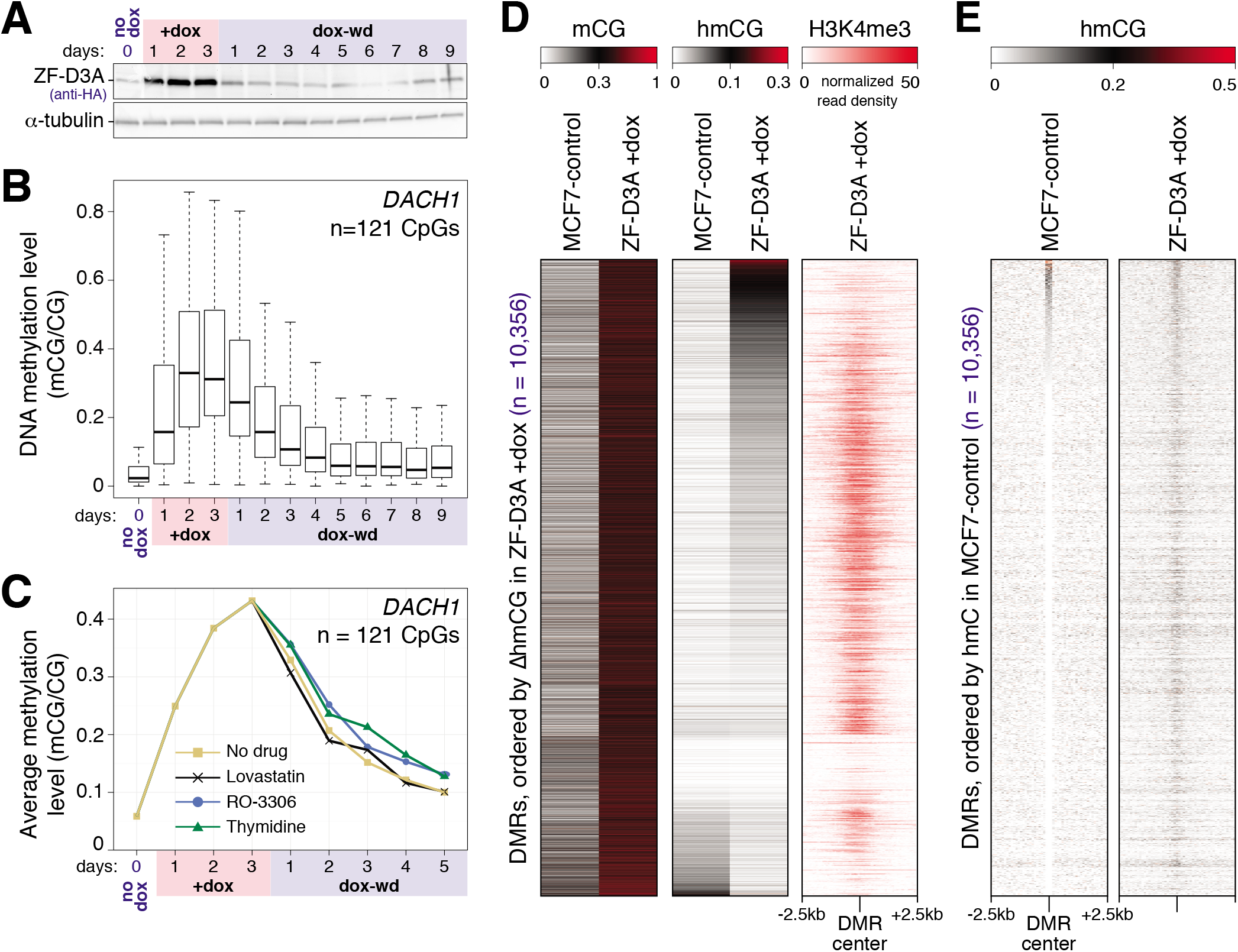
DNA methylation is lost in the absence of ZF-D3A by an active process. **(A)** Western blot measurement of ZF-D3A protein abundance upon doxycycline induction and withdrawal. **(B)** Box and whisker plots of the DNA methylation levels of the 121 CpG sites in the *DACH1* promoter region throughout doxycycline induction and withdrawal. Whiskers indicate 1.5 times the interquartile range or the most extreme data point, whichever is lower. **(C)** Average DNA methylation level at the 121 CpG sites in the *DACH1* promoter region throughout doxycycline induction followed by doxycycline withdrawal and cell cycle inhibition by growth in doxycycline-free media containing different cell cycle inhibitors (Lovastatin, RO-3306, thymidine). **(D)** DMR mCG and hmCG levels in MCF7-control and ZF-D3A +dox cells, ordered by the difference in DMR hmCG level (hmCG/CG) between MCF7-control and ZF-D3A +dox cells, and heatmap representation of H3K4me3 levels in ZF-D3A +dox cells flanking (±2.5 kb, 50 bp bins) the centre of the DMRs. **(E)** Heatmap of hmCG levels flanking (±2.5 kb, 50 bp bins) the centre of the DMRs, ordered by DMR hmCG level in MCF7-control cells.

Targeted bisulfite sequencing analysis of DNA methylation dynamics at several representative promoters exhibiting strongly induced methylation upon ZF-D3A induction showed that the robust increase in DNA methylation under ZF-D3A induction returned to near-baseline levels 5 days after the removal of doxycycline (Fig. 4B, fig. S3 A to F). Thus, both WGBS (Fig. 2, A and C) and targeted bisulfite sequencing analyses demonstrated that forced DNA methylation by ZF-D3A is rapidly removed in the absence of the ZF-D3A protein.

The loss of induced DNA methylation following removal of ZF-D3A could be the result of passive loss by lack of its maintenance after DNA replication, or active removal by a mechanism such as TET dioxygenase-mediated demethylation (*45*). To determine whether DNA methylation loss occurred through an active or passive process, we grew ZF-D3A cells in the presence of doxycycline for 3 days, after which the media was changed to doxycycline-free media containing different cell cycle inhibitors at concentrations previously shown to be effective in MCF7 cells (Lovastatin - G1 block; thymidine - S-phase block) or HeLa cells (RO-3306 - G2/M block) (*46*–*48*). To assess the cell cycle inhibitor efficacies, ZF-D3A cells were treated with each drug for 48 hours and the percentage of cells in G1, S and G2/M phases of the cell cycle quantified. Lovastatin treatment resulted in an increase in the percentage of cells in G1 from 44.4% to 71.5%, while RO-3306 resulted in an increase from 24.2% to 57.1% of cells in G2/M. While thymidine caused a decrease in the percentage of cells in S-phase, a clear shift within S-phase to cells near the S-G2 transition as compared to the untreated cells was observed (fig. S4, A and B). Importantly, despite this disruption of cell cycle progression, these compounds had negligible effect upon DNA methylation loss (Fig. 4C, fig. S4C). For example, after 5 days of doxycycline withdrawal, 88-100% of the induced DNA methylation at the 7 loci analyzed was lost in the cells treated with the three cell cycle inhibitors, compared to the untreated cells, demonstrating that DNA methylation deposited by ZF-D3A is actively removed.

The most thoroughly documented active DNA demethylation mechanism operates via the TET family of dioxygenase proteins, which proceed through a 5-hydroxymethylcytosine (hmC) intermediate (*49*). To assess whether forced methylation was subjected to TET-mediated demethylation, sites of hmC were identified genome-wide by TET-Assisted-Bisulfite-sequencing (TAB-seq) (*50*) in MCF7-control and ZF-D3A +dox cells. A significant increase in hmC levels in the DMRs (p-value <2.2 x 10^−16^, Wilcoxon signed rank test) was observed in ZF-D3A +dox (Fig. 4D, fig. S1B). A significant enrichment in hmC (p-value <2.2 x 10^−16^, Wilcoxon signed rank test) occurred precisely at the DMRs compared to flanking (±2.5 kb) genomic regions in ZF-D3A +dox cells, whereas no significant enrichment was observed (p-value 0.29, Wilcoxon signed rank test) in the same regions in MCF7-control cells (Fig. 4E and fig. S3G). Notably, DMRs with the highest hmC levels were depleted in H3K4me3 in ZF-D3A +dox cells, suggesting that H3K4me3 may stimulate further oxidation of hmC to 5-formylcytosine and 5-carboxylcytosine (Fig. 4D). A small subset of DMRs (12%) exhibited no hmC under any condition, and no H3K4me3 (average read depth <1), suggesting that an hmC-independent process may demethylate some loci. Taken together with the observation that the loss of ZF-D3A induced methylation does not require DNA replication, this strongly implicates TET-mediated active removal of induced methylation at most DMRs.

## Forced promoter methylation is frequently insufficient for transcriptional silencing

Promoter DNA methylation induced by artificial epigenome modifiers fused to DNMT3A were previously reported to downregulate transcription, and have been proposed as therapeutic systems for silencing disease-associated genes (*22*, *23*, *25*). However, the efficacy of targeted DNA methylation-induced silencing and the fraction of genes at which forced DNA methylation is sufficient to induce silencing are unknown. We compared the increase in DNA methylation at UMRs located within 2 kb of transcriptional start sites of expressed genes (>20 mapped reads) with the steady state mRNA levels of the corresponding genes in cells expressing or not expressing ZF-D3A (Fig. 5A, table S4). Of the expressed genes associated with UMRs that are robustly methylated (∆mCG >0.3, n = 2,063) upon ZF-D3A induction, 37% showed either no decrease or a gain in mRNA abundance in ZF-D3A +dox cells (Fig. 5, B [blue columns] and C, fig. S5), while a further 21% exhibited only small decreases (0.7-0.9 fold) in expression (Fig. 5B, green columns) and 42% showed modest to strong repression (Fig. 5B [red columns], mRNA abundance ratio [ZF-D3A +dox / MCF7-control] < 0.7). No correlation was evident between the ability of ZF-D3A to repress transcription and the expression level of the gene (fig. S6, A and B, upper panels), the number of CpG dinucleotides in the UMR (fig. S6, A and B, middle panels), or the CpG dinucleotide density in the UMR (fig. S6, A and B, lower panels). Thus, induced DNA methylation of promoters is frequently insufficient as a primary instructive biochemical signal for gene silencing in these cells.

**Fig. 5.**
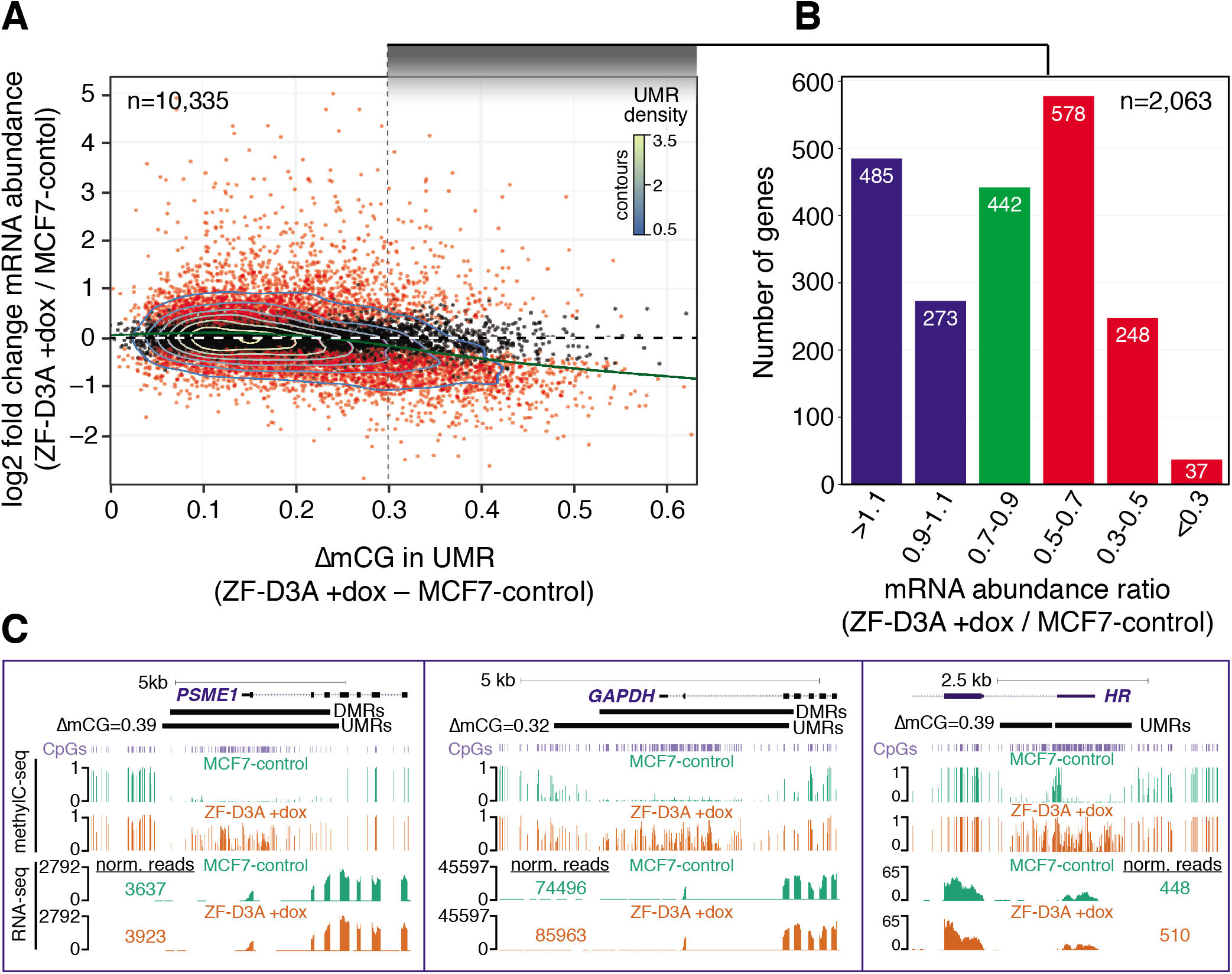
Forced promoter methylation is frequently insufficient for transcriptional repression. **(A)** Scatter plot of the difference in UMR DNA methylation levels, versus the fold change in mRNA abundance of UMR-associated expressed genes, between MCF7-control and ZF-D3A +dox. Point color indicates the gene differential expression significance: red indicates FDR <0.01, black indicates FDR >0.01. Trend line (green) was fitted using a generalized additive model, and contour lines represent the relative density of UMRs. **(B)** Distribution of the ratios of mRNA levels between ZF-D3A +dox and MCF7-control, for expressed genes that have an associated UMR with a ∆mCG >0.3. Column color indicates the class of mRNA abundance ratios between ZF-D3A +dox and MCF7-control: blue = no decrease; green = small decrease; red = decrease. **(C)** Representative genes with high induction of promoter DNA methylation in ZF-D3A +dox and no reduction in mRNA abundance.

Because ZF-D3A does not induce 100% methylation of all CpGs throughout the UMRs, there exists the possibility that two distinct populations of cells are being analyzed: one in which a promoter remains unmethylated and the associated gene is not repressed, and another in which the promoter of the gene is methylated and the gene is silenced. To address this, we performed RNA-fluorescence in-situ hybridization (RNA-FISH) with probes to the *GAPDH* and *PSME1* mRNAs in ZF-D3A +dox cells. These two genes were selected for closer scrutiny because they had UMRs that were strongly methylated by ZF-D3A and showed no reduction in mRNA levels upon ZF-D3A expression (Fig. 5C). ZF-D3A +dox cells were sorted by FACS into high and low expressing subpopulations based on *GAPDH* or *PSME1* RNA-FISH signal. Notably, for both genes the unsorted cells showed a unimodal distribution of target transcript abundance, as opposed to two distinct subpopulations which would be expected if only a subpopulation of cells were gaining DNA methylation (Fig. 6, A and B). FACS reanalysis of the Sorted cells (Fig. 6, A and B), in addition to RT-qPCR (*GAPDH*) or RNA-seq (*PSME1*) (Fig. 6, C and D), demonstrated that RNA-FISH was effective for isolating subpopulations of cells exhibiting high and low levels of the targeted mRNA. Targeted bisulfite sequencing of the *GAPDH* and *PSME1* promoters in the sorted populations revealed that DNA methylation levels were very similar in both low expressing and high expressing cells (Fig. 6, E to G). This further supports our observation that induced DNA methylation of gene promoters is frequently not sufficient to repress transcription.

**Fig. 6.**
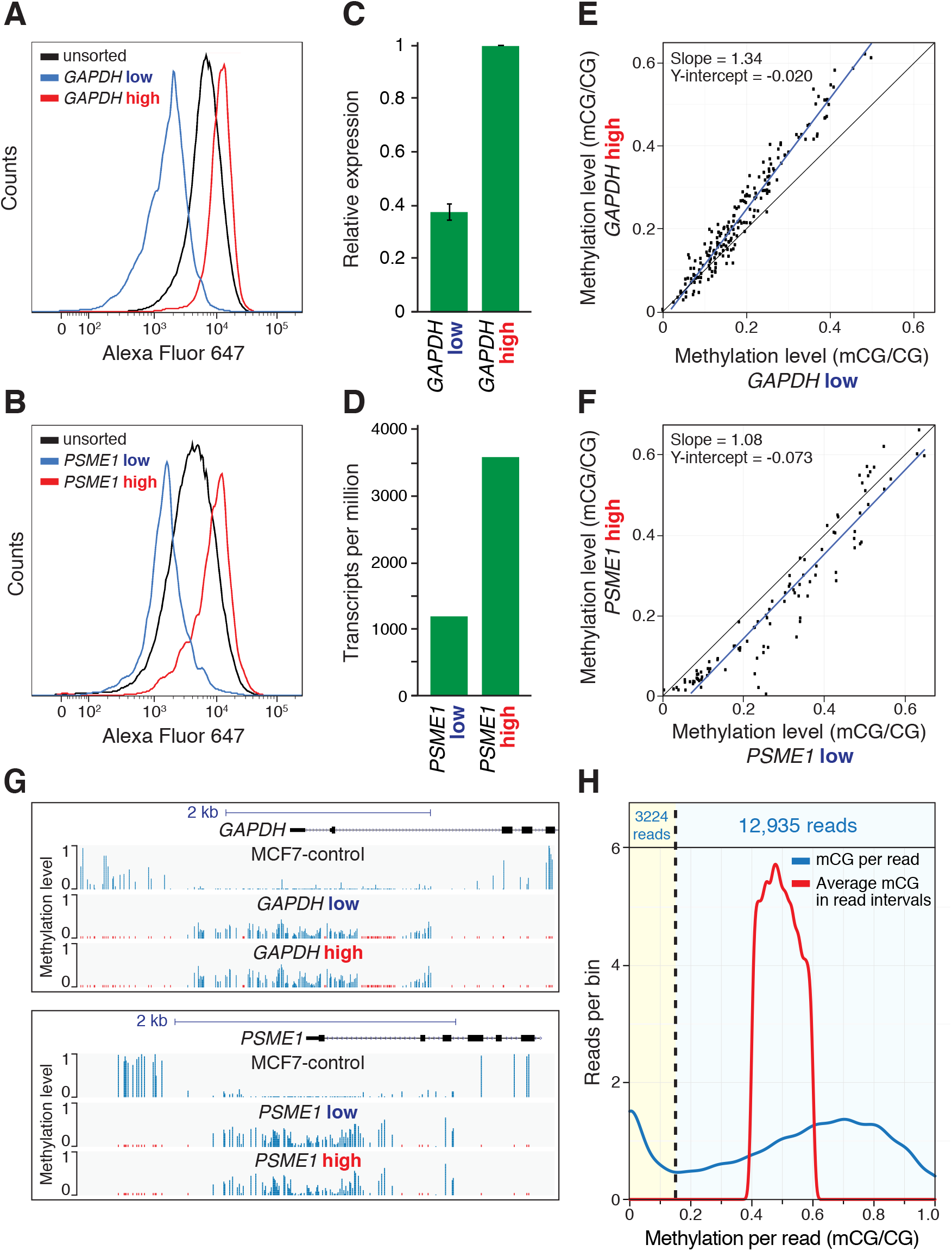
Heterogeneity in gene expression between cells does not reflect differences in DNA methylation. Fluorescence distribution of cells stained with RNA-FISH probes targeting **(A)** *GAPDH* or **(B)** *PSME1* mRNA before and after FACS. mRNA quantitation after FACS sorting of (C) *GAPDH* by RT-qPCR, and **(D)** *PSME1* by RNA-seq. DNA methylation levels of single CpGs at the promoter region of **(E)** *GAPDH*, and **(F)** *PSME1*, for the low (x axis) and high (y axis) expressing sorted cell populations. **(G)** Genome browser representation of the *GAPDH* (top) and *PSME1* (bottom) promoter regions showing DNA methylation in MCF7-control and low and high expressing sorted cell populations. **(H)** Distribution of per-read methylation levels (blue line) for selected ZF-D3A +dox WGBS reads (≥10 cytosines in the CpG context) that align within robustly methylated DMRs (DMR mean methylation level [mCG/CG] of 0.4-0.6 in ZF-D3A +dox and <0.1 in MCF7-control), and average CG methylation levels from ZF-D3A +dox WGBS data of the DMR genomic intervals covered by the selected reads (red line).

To further investigate the possibility that there was a subpopulation of DNA molecules that were not methylated by ZF-D3A, we analyzed the distribution of methylation in single DNA molecules, focussing on CpG-dense WGBS reads (≥10 CpG context cytosines) from ZFD3A +dox cells that aligned to robustly methylated DMRs (DMR mean methylation level [mCG/CG] of 0.4-0.6 in ZF-D3A +dox and <0.1 in MCF7-control). Of 16,159 reads analyzed from ZF-D3A +dox cells, only 20% had a low fraction of methylated cytosines (mCG/CG <0.15, Fig. 6H). In contrast, 80% of reads were methylated modestly to completely (mCG/CG ≥0.15), with 67% of reads exhibiting a robust gain of methylation (mCG/CG ≥0.4, Fig. 6H, blue line). The genomic coordinates to which each read aligned were used to calculate the average methylation in ZF-D3A +dox cells for each read interval (Fig. 6H, red line). While there is a broader distribution of single DNA molecule methylation levels than the selected average range, only a small subset of DNA molecules escape methylation by ZF-D3A, with most being methylated at moderate to high levels. Thus, the inability of ZF-D3A to repress gene expression at a large number of promoters appears to be due to the inability of DNA methylation on its own to induce transcriptional silencing.

## Discussion

DNA methylation is commonly considered a stable and repressive modification, and targeted approaches to manipulate methylation states are being explored for both research and potential clinical applications (*25*). However, very little is known about whether DNA methylation can generally function as a primary instructive biochemical signal for transcriptional silencing of nearby genes and the stability of altered DNA methylation states when induced in the genome. Here we show that the simultaneous methylation of thousands of promoters in the human genome frequently resulted in no detectable repression of gene expression. Induced DNA methylation had very little, if any, effect upon the association of initiated RNA polymerase II or H3K4me3 with DNA, demonstrating that DNA methylation on its own is not sufficient to reconfigure DNA into a stable heterochromatinized state. Finally, forced DNA methylation is rapidly lost at nearly all promoters in the absence of the artificial methyltransferase, likely through TET-mediated demethylation.

It is not entirely clear how the CpG-rich regions associated with transcriptional start sites establish and retain their unmethylated state in a genome that is almost entirely methylated in the CpG context (*5*, *51*). It has been proposed that *de novo* DNA methyltransferases are excluded from these regions (*41*, *52*), however this cannot be the only process, because aberrant methylation deposited in CpG islands would require a proofreading mechanism to prevent the entire genome from drifting towards a fully methylated state. Our observations of induced DNA methylation being actively removed from gene promoters and increased levels of 5-hydroxymethylcytosine in forcibly methylated areas support a model whereby CpG islands are protected from DNA methylation by a TET-mediated surveillance process (*53*, *54*). This is consistent with the observation that the CxxC domains of the TET1 and TET3 proteins specifically bind to CpG dinucleotides with a slight preference for the unmethylated state (*55*–*57*). Furthermore, while ZF-D3A induced large increases in DNA methylation, it did not fully methylate entire promoter regions. The broad distribution of methylation states in individual reads may reflect simultaneous catalysis of methylation by ZF-D3A and demethylation by TET proteins.

Importantly, artificially induced DNA methylation is frequently insufficient for transcriptional repression, even at genes that accumulate high levels of promoter DNA methylation. While DNA methylation is clearly an important modification, there is very little direct evidence that, on its own, it is sufficient to block transcription or force DNA into a heterochromatinized state. Observations that DNA methylation follows transcriptional repression during development have suggested that it acts as a reinforcing mark (*58*). Notably, Amabile *et al.* recently observed that a TALE-DNMT3A fusion protein targeting the *B2M* gene had minimal effects upon transcription in transfected human cells, while simultaneous recruitment of DNMT3A, DNMT3L and KRAB domain proteins resulted in durable and long lasting silencing of the gene (*30*). The KRAB domain recruits several repressors including histone modifying proteins and the nucleosome remodelling complex NuRD The requirement for both the KRAB domain and DNA methylation to stably repress transcription, taken together with our observation that DNA methylation is actively removed when deposited on promoters and the recognition that DNA methylation often follows gene silencing in development, suggests that active chromatin is protected against DNA methylation in part by the TET proteins, and that more extensive changes to multiple DNA and histone modifications may be required simultaneously to affect changes in chromatin structure and gene expression.

This work constitutes the first broad assessment of the repressive capacity of promoter DNA methylation in the human genome. With routine comprehensive epigenome profiling revealing extensive differential methylation between cell types and states, it is now important to establish an extensive understanding of the causal relationships between these covalent modifications and genome regulation. While this study was performed in MCF7 cells, in the future it will be important to undertake comprehensive epigenome manipulation in a wide variety of cell types and states in order to establish the generalisability of these relationships, for example in pluripotent and distinct differentiated cells. Our work demonstrates the utility of artificial epigenome editing technologies for understanding the information content of covalent modifications of the genome, and highlights potential biological barriers that need to be overcome for their use.

## Data access

DNA methylation, TAB-seq, ChIP-seq, and RNA-seq data can be accessed at the Gene Expression Omnibus (GEO) under the accession number GSE102395 and viewed in the UCSC genome browser:

https://genome.ucsc.edu/cgi-bin/hgTracks?hgS_doOtherUser=submit&hgS_otherUserName=ethanford&hgS_otherUserSessionName=FordEtAlBioRxiv2017.

## Acknowledgements

We thank C. Pflüger, M. Cruickshank and M. Lewsey for critical reading of this manuscript, and Andrea Holme at the UWA Centre for Microscopy, Characterisation and Analysis and Ji Kevin Li at the Harry Perkins Institute of Medical Research Flow Cytometry Facility for performing the FACS and flow cytometry analysis. This work was funded by the Australian National Health and Medical Research Council (GNT1069830), the Australian Research Council (ARC) Centre of Excellence program in Plant Energy Biology (CE140100008), the National Institutes of Health (R01CA170370 and 1R01DA036906), the Raine Medical Research Foundation, the National Cancer Institute (P30CA014089), and the National Human Genome Research Institute (R21HG006761). RL was supported by an ARC Future Fellowship (FT120100862), a Sylvia and Charles Viertel Senior Medical Research Fellowship, and a Howard Hughes Medical Institute International Research Scholarship. OB was supported by an Australian Research Council Discovery Early Career Researcher Award (DE140101962). AdM was supported by an EMBO Long Term Fellowship (ALTF144-2014). PB was supported by an ARC Future Fellowship (FT130101767) and by a Cancer Council Western Australia Research fellowship.

## Materials and Methods

### Cell culture

The MCF7-control and ZF-D3A cell lines have been described previously (*25*). Cells were grown in MEM α nucleosides (Life Technologies) supplemented with 0.075% sodium bicarbonate and 10% tetracycline-free fetal bovine serum (Clonetech). Doxycycline induction was performed by adding doxycycline to a final concentration of 100 ng/ml.

### Propidium iodide staining

Ten centimeter cell culture dishes with cells at 30% confluency were grown for 48 hours with or without cell cycle inhibitors and harvested by incubation in PBS with 5 mM EDTA and centrifugation at 900 x g in a swinging bucket rotor. The cell pellet was loosened by vortexing and ice cold 70% ethanol was added dropwise while vortexing and then incubated on ice for 1 hour. The cells were centrifuged at 900 x g for 5 minutes in a swinging bucket rotor and washed twice by resuspending in PBS and centrifuging at 900 x g for 5 minutes. Finally, the cells were resuspended in 100 µl PBS and 5 µl 100 mg/ml RNase A and incubated for 5 minutes at room temperature, after which 100 µl of 100 µg/ml propidium iodide was added and the cells were analyzed on a FACScanto.

### Whole Genome Bisulfite Sequencing by MethylC-seq

Genomic DNA was extracted from cells using the Qiagen Blood and Tissue Kit (Qiagen) according to the manufacturer's instructions. 1 µg of genomic DNA was spiked with 0.5% (w/w) of unmethylated lambda phage DNA (Promega) for calculation of the bisulfite non-conversion rate, and sheared with a Covaris S2 sonicator to an average length of 200 bp. The sheared DNA was end-repaired, A-tailed and ligated to methylated Illumina TruSeq adapters and subjected to 4 cycles of PCR amplification using KAPA HiFi Uracil+ DNA polymerase (KAPA Biosystems). Single-end 100 cycle sequencing was performed on an Illumina HiSeq 1500. Reads were mapped to the human genome (hg19) with the Bowtie alignment algorithm (*60*) as previously reported (*61*).

### TET-assisted bisulfite sequencing (TAB-seq)

Genomic DNA was isolated as described for MethylC-seq. TAB-seq libraries were generated using the 5hmC TAB-seq kit (WiseGene) according to the manufacturer's instructions. 5-hydroxymethylated pUC19 DNA (WiseGene) was used to estimate the protection of 5hmC by β-glucosyltransferase, and unmethylated lambda phage DNA was used to estimate the bisulfite non-conversion rate. Single-end 100 cycle sequencing was performed on a HiSeq1500. Reads were mapped as described for MethylC-seq.

### RNA-seq

RNA was extracted from cells using the Qiagen RNeasy kit. 330 ng of total RNA was used to generate libraries using the Illumina TruSeq Stranded mRNA kit according to the manufacturer's instructions, except one-third of all reaction volumes were used and the final amplification used 10 cycles of PCR. Reads were mapped to the human genome assembly hg19 and transcriptome using TopHat2 (*62*). Uniquely mapped reads were assigned to genes with HTseq-count and differentially expressed genes and fold-change of expression were calculated with DESeq2 (*63*). Normalized read counts from DESeq2 were used to quantitate DNMT3A expression. RNA-seq data was visualized on the UCSC genome browser by normalizing to the sample with the fewest mapped reads. FPKM values were calculated with CuffDiff2 (*64*). For all analyses, only genes with >20 uniquely mapped reads were used.

### ChIP-seq

Chromatin immunoprecipitation followed by massively-parallel DNA sequencing (ChIP-seq) for the HA-tag (for HA-tagged ZF-DNMT3A localization), as well as for the H3K4me3 histone modification, was performed as described previously (*37*).

### ChIP-bisulfite-sequencing

Two 15 cm plates of cells were grown and doxycycline induced for three days. Cells were washed two times with 10 ml of PBS and crosslinked for 5 minutes in 50 mM HEPES-KOH, pH 7.5, 100 mM NaCl, 1mM EDTA, 1% formaldehyde. The crosslinking reaction was quenched by the addition of glycine to a final concentration of 125 mM and washed twice with phosphate buffered saline (PBS). All subsequent solutions were supplemented with a protease inhibitor cocktail (Sigma Cat. # P8340). Cells were scraped off the plates with a rubber policeman in 10 ml of PBS and centrifuged at 3,000 rpm for 5 minutes in a swinging bucket rotor. The cell pellets were resuspended in 10 ml of 50 mM HEPES-KOH, pH 7.9, 140 mM NaCl, 1 mM EDTA, 10% glycerol, 0.5% NP-40, 0.25% Triton X-100, incubated on ice for 10 minutes and centrifuged at 3,000 rpm for 10 minutes in a swinging bucket rotor. Cell pellets were washed twice by gently adding 10 mM Tris-HCl, pH 8.1, 200 mM NaCl, 1 mM EDTA to the cell pellets, trying not to disturb the pellets, and centrifuged at 3,000 rpm for 5 min. Finally the cell pellets were resuspended in 0.1% SDS, 1 mM EDTA and transferred to a Covaris TC12x12 tube. The chromatin was sheared using a Covaris S2 sonicator with the following settings: time 12 min, duty cycle 5%, intensity 4, cycles per burst 200, temperature 4˚C, power mode frequency sweeping. Triton X-100 and NaCl were added to a final concentration of 1% and 150 mM respectively. The sheared chromatin was centrifuged at maximum speed in a microfuge for 15 minutes at 4˚C and the supernatant was transferred to a new tube. 2 µl of anti-H3K4me3 (Diagenode, Cat. # C15410003) or 4 µl of anti-phospho-Ser5 RNA polymerase II antibody (Active Motif, Cat. # 39233) was added and incubated overnight at 4˚C. 30 µl of Protein G Dynabeads (Life Technologies) was added and incubated on a tube rotator for 90 minutes at 4˚C. The beads were washed twice with 20 mM HEPES-KOH, pH 7.9, 0.1% SDS, 150 mM NaCl, 1% Triton X-100 2 mM EDTA, twice with 20 mM HEPES-KOH, pH 7.9, 0.1% SDS, 500 mM NaCl, 1% Triton X-100 2 mM EDTA, once with 100 mM Tris-HCl pH 7.5, 0.5 M LiCl, 1% NP-40, 1% sodium deoxycholate, and once with 10 mM Tris-HCl, pH 8.0, 1 mM EDTA. The DNA was eluted twice by incubating for 30 minutes in 25 µl of 20 mM HEPES-KOH, pH 7.9, 1 mM EDTA, 0.5% SDS, 0.5 mg/ml Proteinase K. To the 50 µl of eluted DNA, 3 µl of 3M sodium acetate, pH 5.3 and 0.5 µl 30 mg/ml RNase A was added and incubated overnight at 65˚C in a hybridization oven. 1.5 µl of 20 mg/ml proteinase K was added and incubated for 1 hour at 50˚C and the DNA was purified with 2 volumes of SPRI beads and eluted in 20 µl Tris-HCl, pH 8.0, 0.1 mM EDTA. Libraries were made with the Accel-NGS Methyl-Seq DNA Library Kit (Swift Biosciences) according to the manufacturer's instructions. Reads were aligned and DNA methylation sites were identified as described for MethylC-seq above, except after adapter trimming 10 nt were hard cropped off the 3' end of the read.

### RNA fluorescence-in-situ-hybridization and FACS sorting

Cells were labeled with fluorescent probes to *GAPDH* mRNA or *PSME1* mRNA using the PrimeFlow RNA kit (eBiosciences) according to the manufacturer's instructions. Cells were sorted into high and low expression populations on a FACS Aria2 by the Harry Perkins Cell Sorting Facility. The sorted cell populations were aliquoted into two tubes for RNA and DNA extraction.

### RNA isolation and quantitation from FACS-sorted cells

RNA was extracted using the Qiagen RNeasy FFPE kit according the manufacturer's instructions. Exon spanning primers were designed to the *GAPDH* and *RPL13A* as an internal control. RNA was reverse transcribed with gene specific primers using SuperScript II (Life Technologies) and qPCR was performed on an LC480 thermocycler using KAPA SYBR FAST

DNA qPCR Master Mix (Kapa Biosystems). For the *PSME1* high and low expressing population, the RNA abundance was quantitated by RNA-seq. Ribosomal RNA was depleted from the sample using the RiboCop rRNA Depletion Kit (Lexogen) and libraries were generated with the TruSeq Stranded mRNA Kit (Illumina). The libraries were sequenced on an Illumina HiSeq 1500 with single-end 100 cycle sequencing. The reads were adapter and quality trimmed with cutadapt (*65*) and transcripts per million were calculated using Kallisto (*66*).

### Targeted Bisulfite-PCR amplicon sequencing

DNA was extracted from FACS sorted cells and bisulfite converted with the EZ DNA Methylation-Direct Kit (Zymo Research). PCR amplicons to the *GAPDH* and *PSME1* promoters were designed with methprimer and genomic DNA was amplified with 40 cycles of PCR using EpiMark Hot Start Taq DNA Polymerase (New England Biolabs). PCR reactions were pooled and purified with a homemade version of AMPureXP beads. 1 µg of pooled PCR products in 14.5 µl were phosphorylated by adding 15 µl 2X Quick Ligase Buffer (New England Biolabs) and 0.5 µl T4 polynucleotide kinase (New England Biolabs) and incubating at 37˚C for 30 minutes. Illumina TruSeq adapters synthesized by IDT and annealed by heating to 99˚C and slowly cooling to 20˚C were ligated to the phosphorylated PCR products by adding 3.75 µl 10 µM annealed TruSeq adapters, 10 µl 2X Quick Ligase Buffer (New England Biolabs) and 6 µl water. The ligation reactions were incubated at 25˚C for 20 minutes and stopped by adding 2 µl 0.5 M EDTA. The DNA was purified by adding 20.8 µl (0.4 volumes) of SPRI beads. The libraries were subjected to single-end 300 cycle sequencing on the Illumina MiSeq. The reads were adapter trimmed with cutadapt and aligned to the human genome (hg19) with BS-Seeker2 (*67*) and bowtie2 (*68*). Methylation was called using bs-seeker2.

### DNA methylation data analysis

DMRs between MCF7-control cells and ZF-D3A +dox cells were called using the software package DSS (*69*). DSS was run with a low p-value cutoff to obtain the DMR boundaries with the following parameters: smoothing=TRUE, smoothing.span=500, delta=0.1, p.threshold=0.05, minCG=4, dis.merge=1000, minlen=100, pct.sig=0. To obtain a more stringent set of DMRs, DSS was run a second time with the same parameters, except the p-value threshold was decreased to 0.005. DMRs from the less stringent set that were also found in the more stringent set were used as the final set of DMRs for subsequent analyses.

UMRs were identified by combining the data sets from MCF7-control, MCF7-control/doxycycline withdrawn and ZF-D3A no-dox cells and were called using the software package methylSeekR (*39*) with the FDR cutoff set to 0.05. To filter UMRs that were called in large hypomethylated regions, UMRs that were within 40 kb were merged into single genomic intervals. The resulting set of genomic intervals were filtered for regions >20kb and had an average methylation of mCG/CG <0.3. UMRs form the original set that did not overlap with the UMRs from large unmethylated regions were used for further analyses.

The weighted DNA methylation level in DMRs and UMRs were calculated by dividing the sum of C base calls by the sum of C + T base calls at all CG dinucleotide positions in the reference genome in the interval. Methylation heatmaps were made by dividing DMRs into 100 bp bins and calculating sum of C base calls in the genomic interval by the sum of C + T base calls in each bin. The resulting matrix was plotted with MeV (*70*). Smooth scatter plots were made with the smooth.scatter R function. All other plots were made with ggplot2 in R. Replicate methylomes were compared by selecting DMRs that had ≥20 base calls in all samples and plotting the resulting heatmap with the R function p.heatmap using the default parameters.

UMRs were assigned to genes by calculating the distance of the UMRs to the nearest transcriptional start site annotated by UCSC (*71*) with the closestBed function of the BEDTools software package (*72*). UMRs that fell within 2 kb of a transcriptional start site were assigned to the respective gene. Genes with ≤20 mapped RNA-seq reads were excluded from the analysis.

Intersection of DMRs and UMRs was performed using the pybedtools functions venn_gchart and venn_mpl. UMRs with multiple intersections with DMRs were counted multiple times. To compare the methylation levels in UMRs with the retention of H3K4me3, UMRs were filtered for those that contained >100 mC base calls and >100 H3K4me3 ChIP-seq reads mapped to the UMR.

The analysis of single WGBS reads (Fig. 6H) was performed by sub-selecting DMRs with a ∆mCG between 0.4 and 0.6, a mCG level of <0.1 in MCF7-control cells, and ≥50 mapped reads. Reads mapped to DMRs that extended beyond the DMR border were trimmed to the DMR interval. The resulting reads mapped to DMRs were sub-selected for reads with ≥10 cytosines in the CpG base context and for intervals that had average methylation levels between 0.4 and 0.6 in ZF-D3A +dox cells. The number of C and T base calls in each read was calculated in Python and plotted with ggplot2 in R.

### ChIP-seq and ChIP-bisulfite-sequencing data analysis

For ChIP-seq, reads were mapped with the bowtie software package (*60*) and non-uniquely mapped reads were discarded. For ChIP-bisulfite-seq, reads were mapped and methylation was called as described for MethylC-seq. Peaks were called using MACS1.4 (*74*). H3K4me3 peaks from ZF-D3A no-dox and ZF-D3A +dox samples were combined and overlapping peaks were merged to the outer boundaries of each peak. Heatmaps were generated by dividing DMRs into bins of 50 bp and counting the reads in each bin. The resulting matrices were plotted with MeV (*70*). ChIP-seq reads for H3K4me1 and H3K27ac in MCF-7 cells were obtained from the GEO database (GSE38447) (*75*).

H3K4me3 ChIP-seq data sets for ZF-D3A +dox and ZF-D3A no-dox cells were normalized by identifying common peaks between the two samples. Peak intervals with an average mC level of <0.2 were sub-selected and the number of reads mapped to ±50 bp of the peak center was used as the normalization factor. Normalized mapped reads counts in H3K4me3 read ZF-D3A no-dox and ZF-D3A +dox cells was measured across an interval ±2.5 kb of the center of DMRs. The resulting intervals with more than 2,000 reads summed from both samples (n=2,735) were used to compare H3K4me3 occupancy between ZF-D3A no-dox and ZF-D3A +dox cells.

Heatmaps of H3K4me3 read density were made using the mapBed function of BEDtools and in R using the p.heatmap function. H3K4me3 reads ±100 bp of the center of DMRs were calculated with the mapBed function of BEDtools and plotted with ggplot2 in R. H3K4me3 reads mapped to UMRs were also calculated with the mapBed function of BEDtools. UMRs with >100 mapped H3K4me3 ChIP-seq reads and >100 cytosine base calls at cytosines were plotted with ggplot2 in R.

---

**Supplementary Fig. S1.**
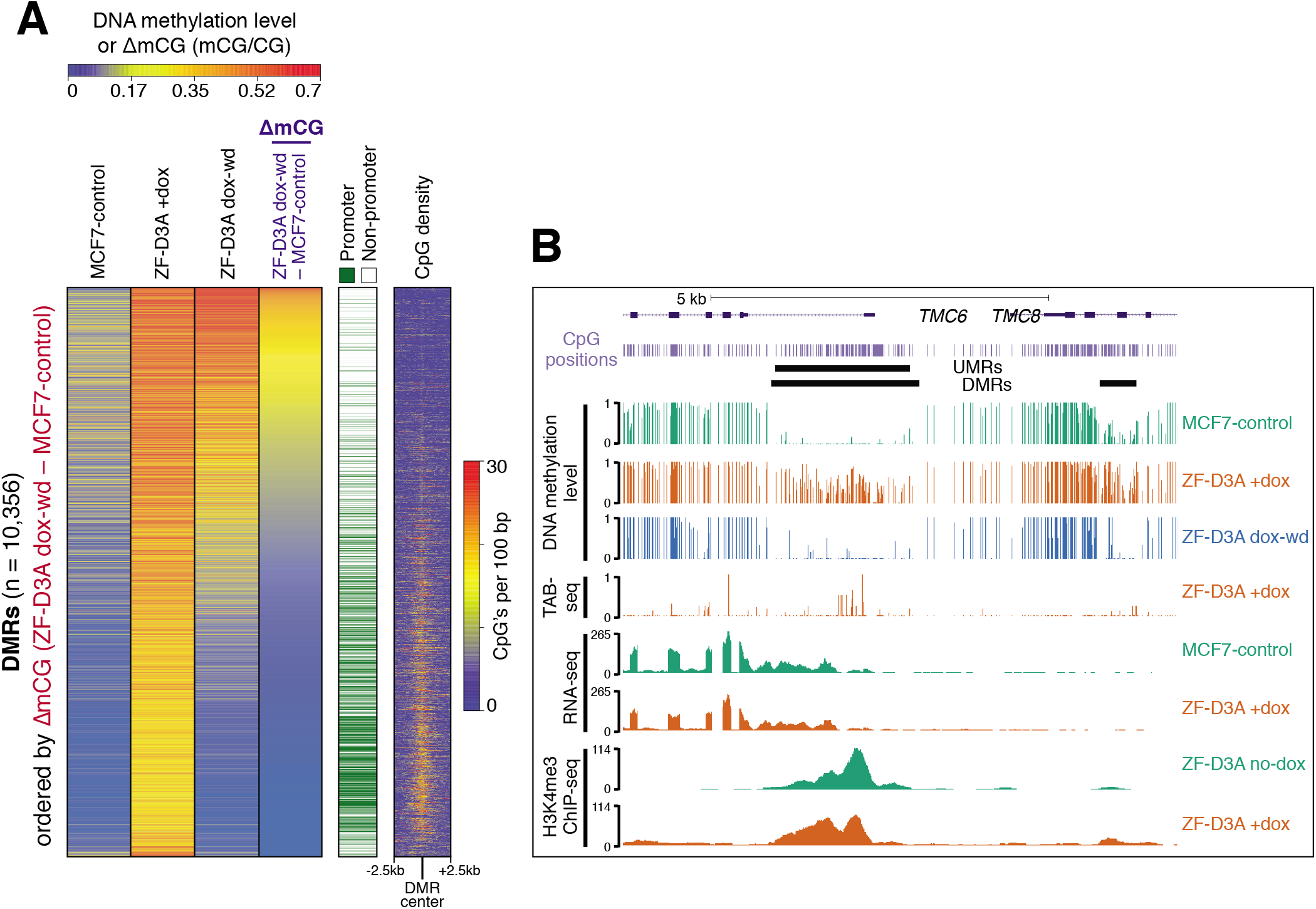
**(A)** Heatmap of DNA methylation levels and differences in all DMRs (n = 10,356) ordered by decreasing ∆mCG (ZF-D3A dox-wd – MCF7-control) in DMRs, classification of each DMR as promoter or non-promoter located, and CpG dinucleotide density flanking (±2.5 kb, 100 bp bins) the center of all DMRs. **(B)** Genome browser screenshots of a representative locus showing off-target DNA methylation by ZF-D3A in the *TMC6* and *TMC8* promoter regions.

**Supplementary Fig. S2.**
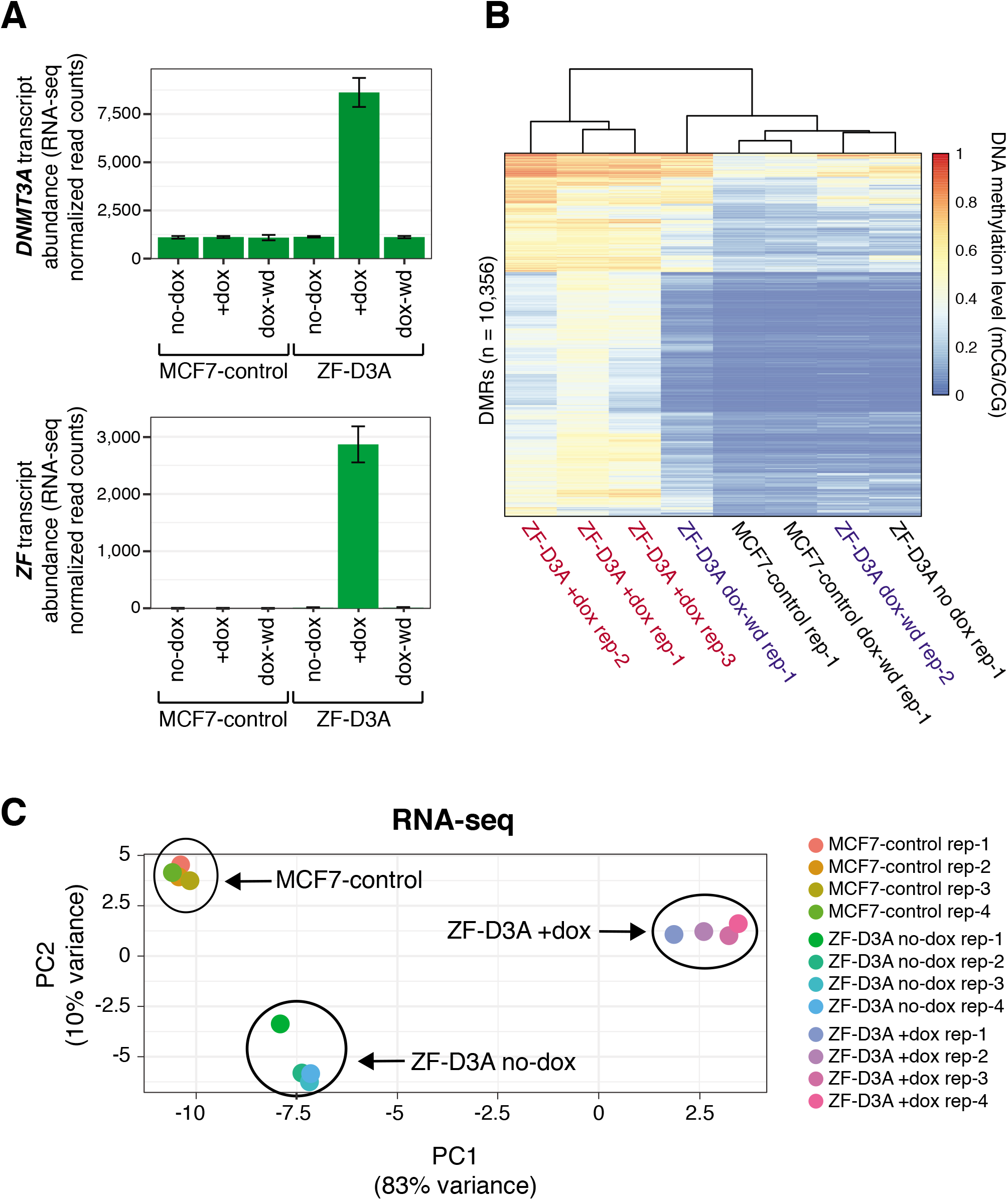
**(A)** Quantitation of *ZF* and *DNMT3A* transcript abundance by RNA-seq in MCF7-control and ZF-D3A cells grown either with no doxycycline (no-dox), in the presence of doxycycline (3 days; +dox), or after doxycycline withdrawal (9 days; dox-wd). Error bars represent standard error of the mean. **(B)** DNA methylation levels (mCG/CG) in DMRs in all WGBS datasets. (C) PCA plot of biological replicate RNA-seq datasets.

**Supplementary Fig. S3.**
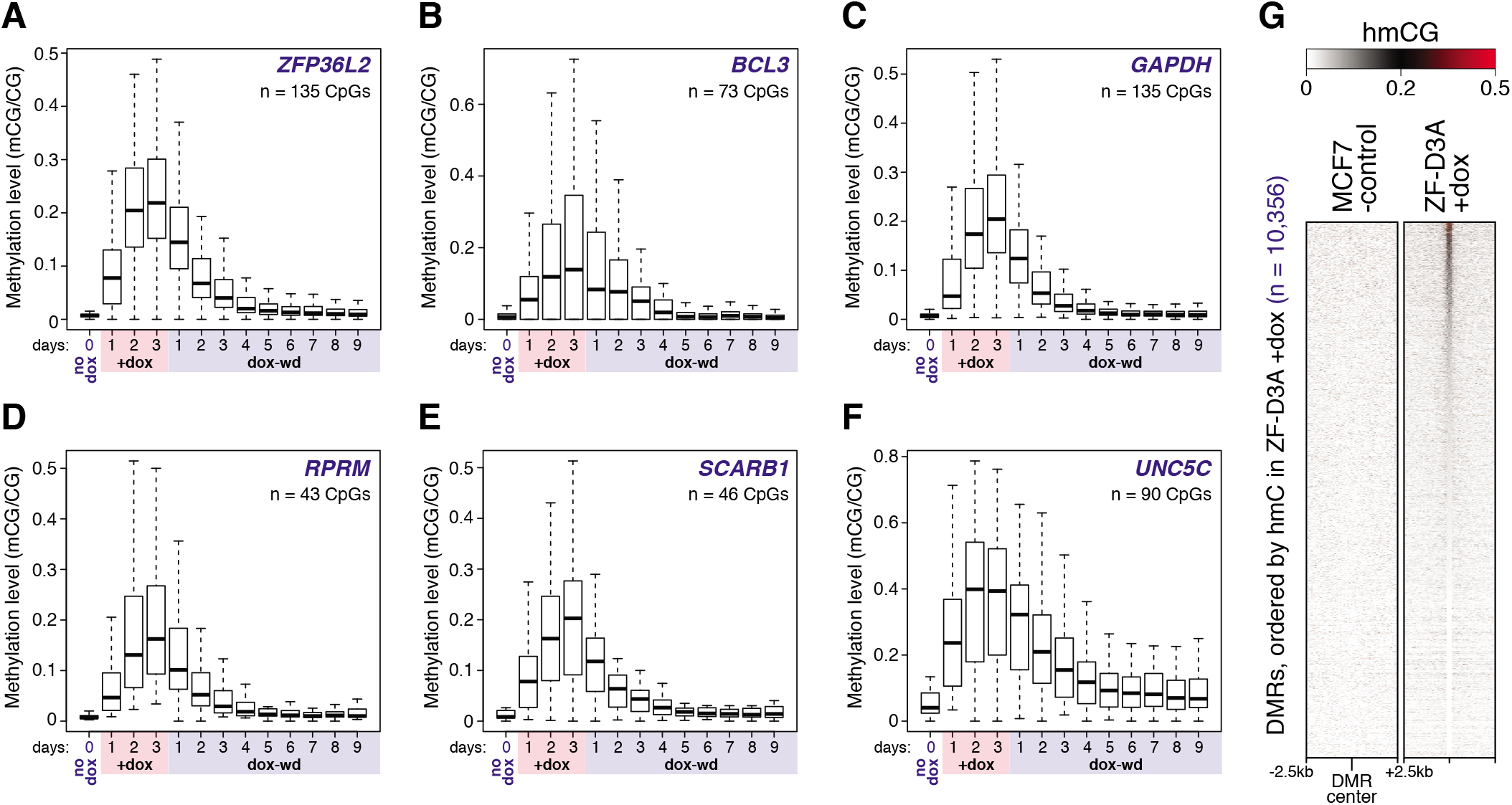
**(A to F)** Box and whisker plots of the DNA methylation levels of the CpG sites in the promoter regions of six selected genes throughout doxycycline induction and withdrawal in the ZF-D3A cell line. Whiskers indicate 1.5 times the interquartile range or the most extreme data point, whichever is lower. **(G)** Heatmap of hmCG levels flanking (±2.5 kb, 50 bp bins) the centre of the DMRs, ordered by DMR hmCG level in ZF-D3A +dox cells.

**Supplementary Fig. S4.**
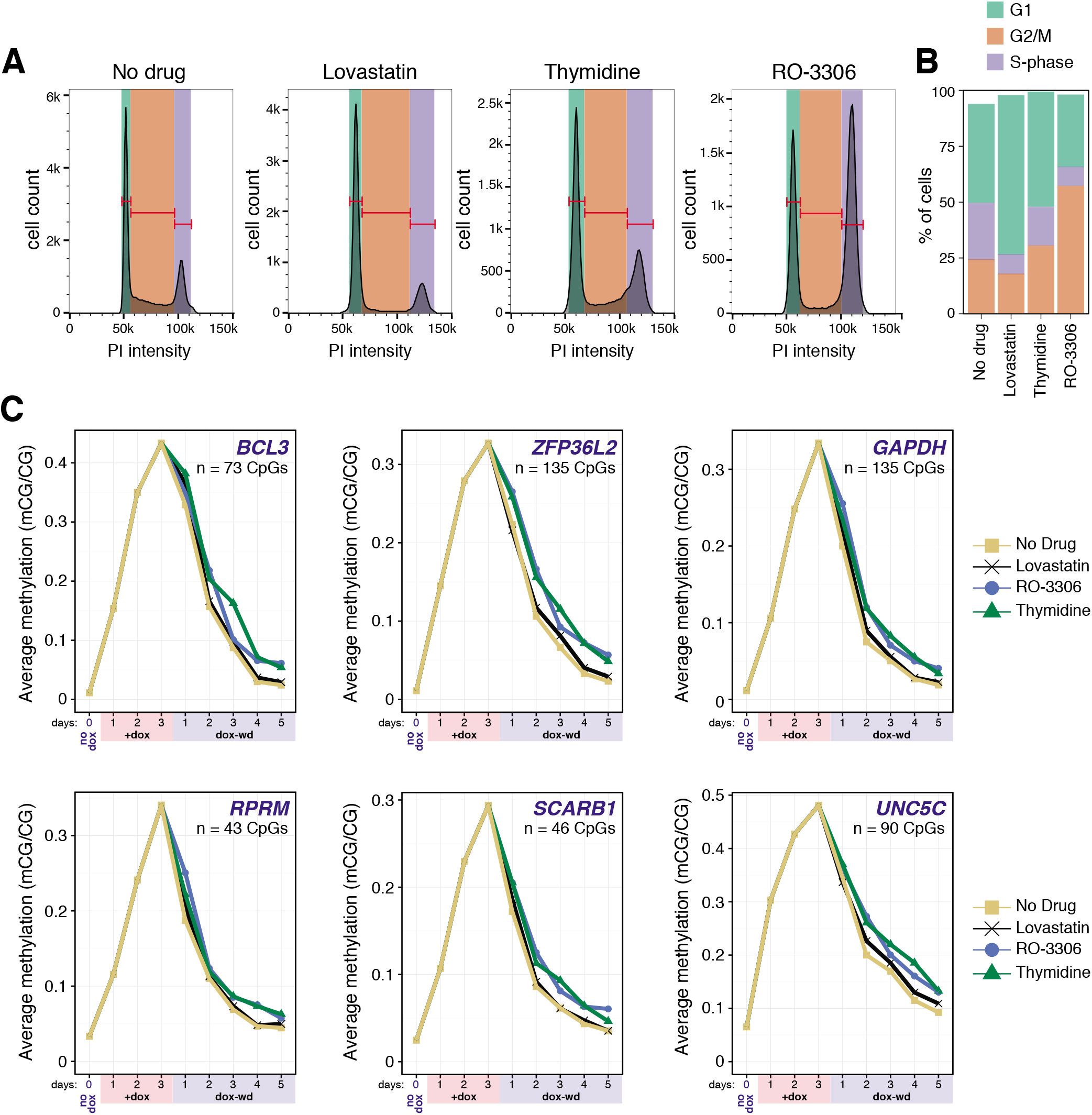
**(A)** FACS analysis of propidium iodide stained MCF7-ZF-D3A cells that had been grown with and without treatment with cell cycle inhibitors. **(B)** Cell cycle distribution before and after cycle inhibitor treatment. **(C)** Average DNA methylation level at the CpG sites in the promoter regions of six selected genes throughout doxycycline induction followed by doxycycline withdrawal and cell cycle inhibition by growth in doxycycline-free media containing different cell cycle inhibitors (Lovastatin - G1 block; thymidine - S-phase block; RO-3306 - G2/M block) and an untreated (no drug) control.

**Supplementary Fig. S5.**
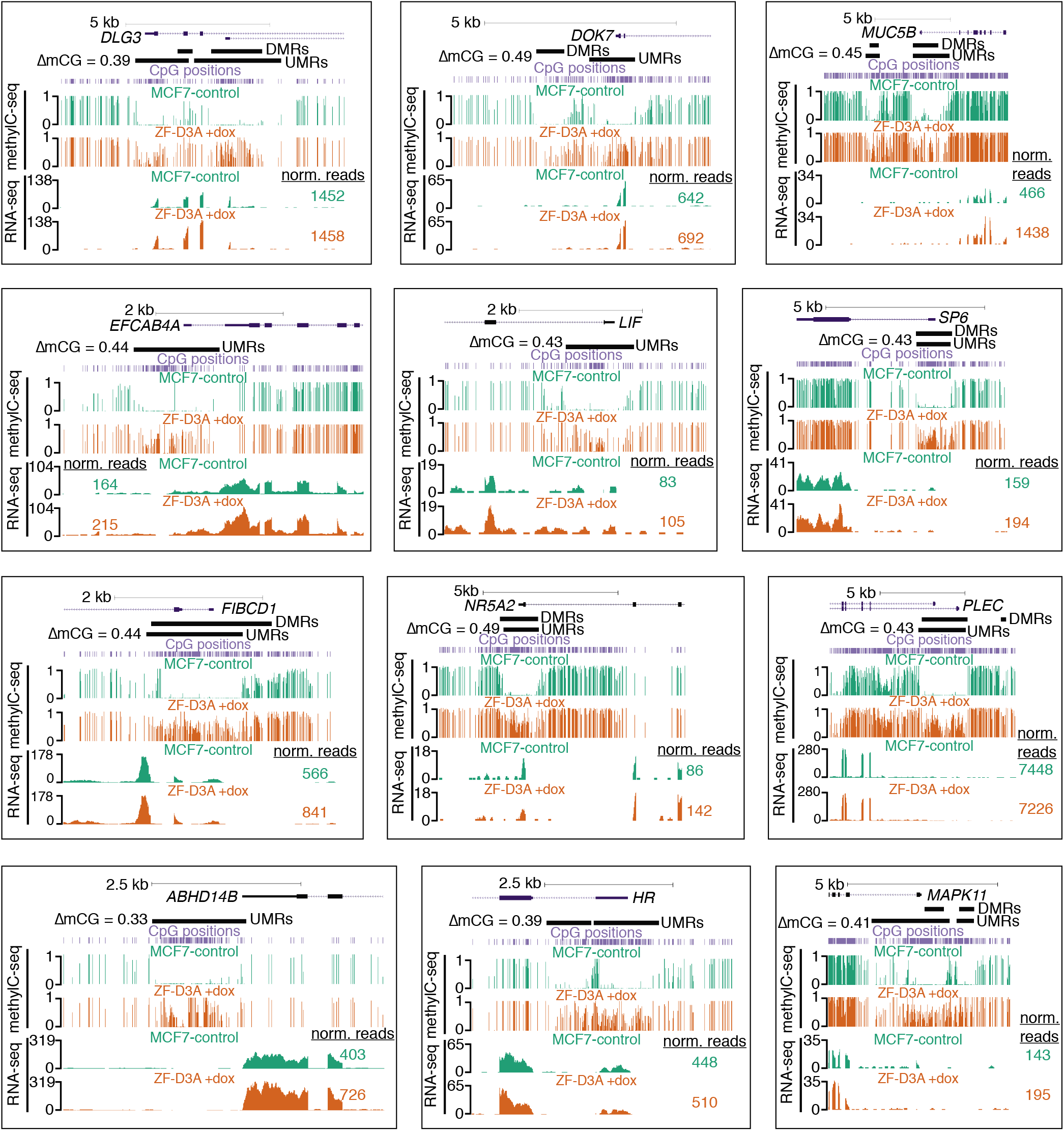
Genome browser screenshots showing DNA methylation sites and levels and RNA expression signal (RNA-seq) for genes with high induction of promoter DNA methylation in ZF-D3A +dox and no reduction in mRNA abundance.

**SupplementaryFig. S6.**
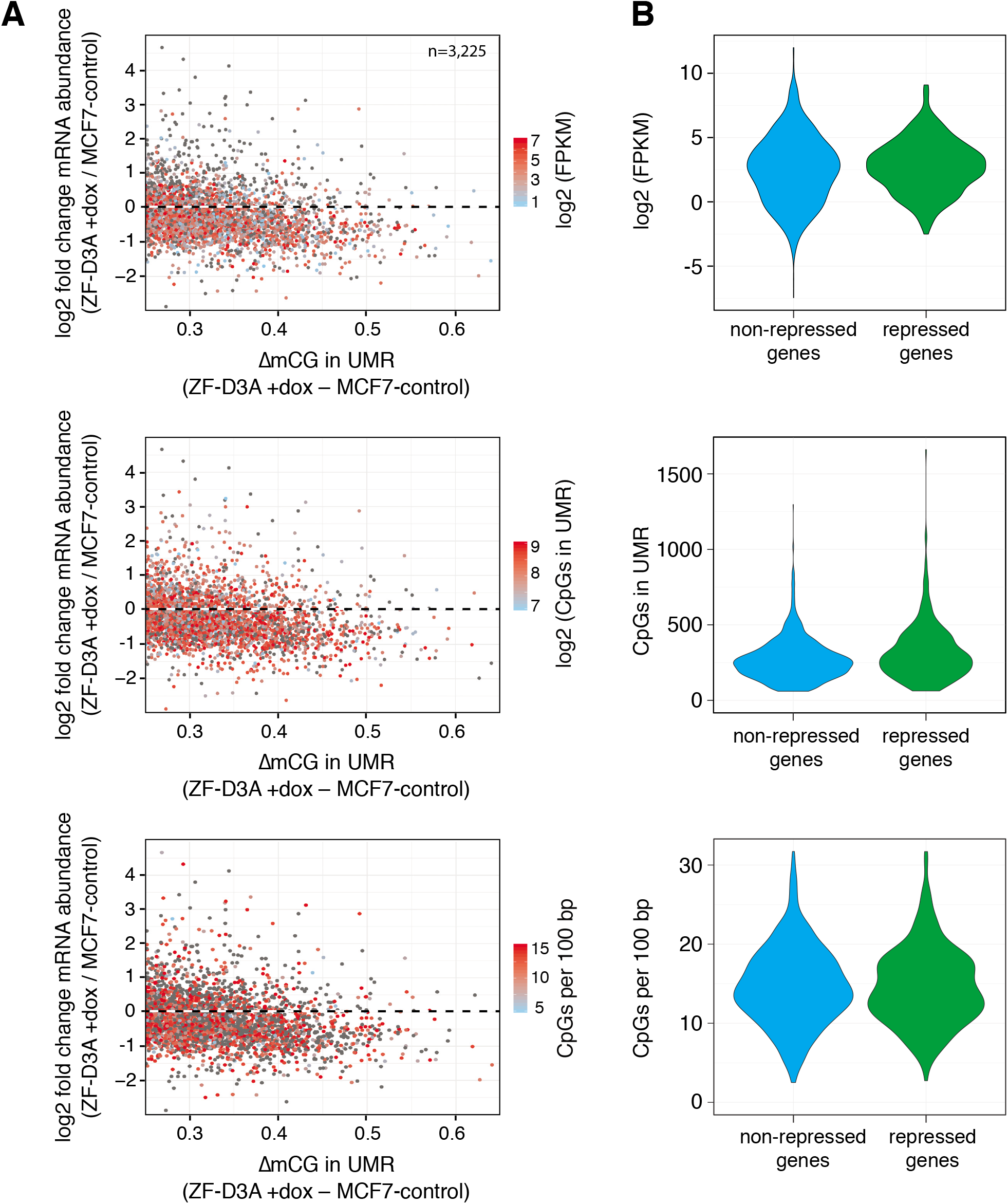
**(A)** Scatter plot of the difference in UMR DNA methylation levels, versus the fold change in mRNA abundance of UMR-associated expressed genes, between MCF7-control and ZF-D3A +dox. Point color indicates relevant UMR feature quantitation. Repression of transcription following induction of ZF-D3A does not correlate with gene expression level (top), number of CpG dinucleotides in the UMR (middle), or CpG density in the UMR (bottom). **(B)** UMRs with a ∆mCG >0.25 were divided into repressed and non-repressed genes. Genes in the repressed and non-repressed categories were required to have a log2 fold change of less than -1 or greater than -0.1 mRNA abundance between the MCF7-control and ZF-D3A +dox data sets, respectively.

